# Effects of mineral addition on the establishment of grazing lawns in a nutrient poor savanna

**DOI:** 10.1101/2024.01.20.576489

**Authors:** Bradley Schroder, Frank Van Langevelde, Nicola-Anne Hawkins Schroder, Herbert H. T. Prins

## Abstract

Nutrient poor savannas are often characterized by inedible or rarely palatable grasses, which generally provide poor nutrition for mammalian grazers. So-called grazing lawns, with short, stoloniferous edible grasses, could provide high-quality food for grazers, but these lawn grasses are rare in nutrient poor savannas. We tested whether we could use mineral addition to establish grazing lawns in a nutrient poor African savanna, in order to achieve a switch from tall, nutritionally poor to short, highly nutritional grass species. The key finding is that phosphorus and lime, nitrogen and nitrogen and lime supplementation resulted in shift from tall to short grasses within three years, with a higher overall nutrient concentration in the grass leaf, than without supplementation. When grazed, the cover of lawn grasses was higher compared to the other grasses when not grazed, demonstrating the role of grazers in maintaining and expanding lawn grass patches. We conclude that local fertilisation in nutrient poor savannas is a viable method of increasing mineral levels in the soil and grass leaf. We also concluded that grazing results in an increase in lawn grass cover and a combination of fertilisation and grazing can improve forage quality to ensure higher nutrient availability to herbivores.

## Introduction

In nutrient poor savannas, tall bunch grass is the predominant grass type, which generally have a low nutritional value that can only support small numbers of mammalian grazers (Bond and Archibald 2003; Parker 2004; Mucina and Rutherford 2006). In these areas, fires are frequent due to high fuel load and thus the so-called grazing lawns are nearly always absent (Bond and Archibald 2003). Grazing lawns are areas with nutritious, grazing-tolerant grass species (lawn grasses) with a short-stature, such as couch grass (*Cynodon dactylon;* a.k.a. Bermuda grass). Lawn grass patches are frequently interspersed with patches of tall bunch grasses, which often have high stem to leaf ratio and high C:N concentrations, and can therefore be less digestible than species found on grazing lawns (Prins and Beekman 1989; Prins 1996; Augustine et al. 2019). The tall bunch grasses grow in clusters and are intolerant to both drought and herbivory, such as common thatching grass (*Hyparrhenia hirta*) (Van der Plas et al. 2013; Archibald et al. 2005; Archibald 2008; Coetsee et al. 2010). Grazing lawns are established through frequent and intense grazing (Grant and Scholes 2006; Arnold 2012; Hempson et al. 2014), and vegetation is observed to switch from lawn grasses to tall bunch grasses in the absence of grazing (O’Connor 1994; Uher-Koch et al. 2019). There are few studies which show the mechanisms for the formation of grazing lawns and what induces the grazing lawns to establish and be maintained. If grazing pressure is relaxed, the short statured lawn grasses can be shaded out and will be outcompeted by the invasion of tall grasses; with frequent fires tall grasses have a rapid post-fire regrowth, which provides little opportunity for other grasses to establish (Cromsigt and Olff 2008; Novellie and Gaylard 2013; Hempson et al. 2014; Hempson et al. 2019).

Grazing lawns are generally found in areas with relatively high soil nutrient status and high grazing pressure (Vesey-FitzGerald 1974; Loth and Prins 1986; Grant et al. 2011; Seymour et al. 2013; Veldhuis et al. 2014). The high soil mineral concentrations potentially lead to higher mineral concentrations in the plants. Owing to these high mineral values, mammalian herbivores will concentrate their grazing in these areas. The more the area is utilised by grazers, the quicker the switch from tall unpalatable to short palatable grasses (Gillard 1969; Coetsee et al. 2010; Novellie and Gaylard 2013; Hempson et al. 2014). Hippopotamus (*Hippopotamus amphibius*) and square-lipped rhinoceros (*Ceratotherium simum*) are examples of grazers which may create and maintain grazing lawns (Vesey-Fitzgerald 1974; Owen-Smith 1988; Cromsigt and Olff 2006; Verweij et al. 2006; Grant et al. 2011).

Grazing lawns are often found on old (abandoned) pastural lands (Valls Fox et al. 2015) or former cattle kraals, where the collective cattle manuring over a long period of time has led to locally high soil nutrient status (Van der Waal et al. 2011a, b; Augustine et al. 2011). This is more prevalent in areas with low annual rainfall, which reduces leaching and results in nutrient hotspots persisting for decades (Young et al. 1995; Veblen 2012). Grazing lawns do not carry fire due to the low fuel load (Bond and Archibald 2003) and can typically be described as a herbivore-driven system. Such a herbivore-driven system is stimulated by grazers, which create heterogeneity in the landscape, with patches often grazed and re-grazed (Prins 1996; De Knegt et al. 2008). In contrast, a fire-driven system generally has a homogenising influence on a landscape (Archibald and Bond 2004; Groen et al. 2017). Van Langevelde et al. (2003) and Bond and Archibald (2003) suggest that fire can indirectly reduce grazing lawns due to creating green flushes in other areas, which draw the grazers away from the lawns, resulting in less grazing pressure and hence a potential increase of tall grasses at the expense of lawn grass species. In areas with higher fire frequencies, grazing lawns seem to be infrequent (Bond and Archibald 2003; Thapa et al. 2021).

There is a large amount of evidence to prove that nutrient addition, either through dung/urine deposition in livestock corrals (Augustine et al. 2011; Van der Waal et al. 2011; Porensky and Veblen 2015), termite mounds (Gosling et al. 2012), or patches of sodic soils (Grant and Scholes 2006) stimulates nutrient hotspots. It remains unknown whether local fertilisation in a nutrient poor system may trigger a transition from tall grass species to short grass species (Coetsee et al. 2010; Zwerts et al. 2015), however it has been proven that elevated soil nutrient levels attract mammalian herbivores to graze, especially when N is added (Schroder 2021; Thapa et al. 2021). Experiments have been undertaken with the combination of N, P and K being simultaneously added to mowed plots (Cromsigt and Olff 2008), which showed that grazing lawns developed in the fertilised areas but not in the unfertilised areas. We tested whether we could use fertilisation with nitrogen, phosphorus and with a combination of lime (calcitic and dolomitic lime) as a possible mechanism by which grazing lawns could be established in Welgevonden Game Reserve, a nutrient poor savanna in the north of South Africa. Our hypothesis is that the addition of minerals (N, P and a combination with Ca) (i.e., artificial fertilisation) into a nutrient poor savanna system will increase the minerals in the soil and thus have a positive effect on the mineral levels in the grass leaf (that is, the available grass for the herbivore species). We further predict that in conjunction with grazing pressure there will be a decrease in tall less palatable grasses and an increase in the abundance of lawn grasses such as couch grass in that study area. The Welgevonden grass species list with the grazing values is available in appendix A. The minerals utilised as artificial fertiliser were chosen due to the limitation of this minerals in nutrient poor savannas and the importance of both N and P to grass growth and survival. N is used for the production of chlorophyll for photosynthesis as well as being a major component in the building of amino acids, proteins and DNA (Martin 2017), whilst P is required by plants to produce Adenosine triphosphate (ATP), the energy carrying molecule found in all living things for cell division and development of new tissue. It should be noted that a deficiency of P has a potential negative effect on the growth and lactation for grazing animals (Murry 1995; Ludwig, de Kroon & Prins 2008) and thus increased levels of these minerals in grass leaf in areas with limited P availability could potentially attract grazers. Several studies have shown that savannas are either limited by N or P (Augustine et al. 2003; Cech et al. 2008). In a study close to Kruger National Park, South Africa, Van der Waal et al. (2011a) found that P fertilisation increased leaf P concentrations in grasses on nutrient poor granite-gneiss soils, more than the expected response of grasses to N fertilisation. This study aims to provide evidence that the addition of minerals and increased grazing in a nutrient poor savanna system can produce short, highly nutritional grass species and give them the opportunity to outcompete the tall, nutritionally poor grass species, establishing and maintaining productive grazing lawns.

## Methods

The study was conducted in Welgevonden Game Reserve, situated on the Waterberg Plateau in South Africa (24°10’S; 27°45’E to 24°25’S; 27°56’E). The average rainfall in the study area is 650 mm per annum (annual rainfall for the study area - supporting documentation A), with most of the area characterized primarily by the soil types dystrophic to mesotrophic yellow-brown apedal coarse sands (Parker 2004). Welgevonden is characterised by ferruginous soils with a low pH (because it is derived from ancient sandstone with high pyrite content; *cf*. Bester and Vermeulen 2010; Burne 2015).

Accordingly, the vegetation type is dominated by broadleaved savanna with tall, nutritionally poor grasses such as *Hyparrhenia hirta* (common thatching grass). The area has a diverse number of grazers, which is dominated by Burchell’s zebra (*Equus quagga burchellii*), blue wildebeest (*Connochaetes taurinus*) and square-lipped rhinoceros (herbivore species list for the study area - supporting documentation B). We tested our hypotheses in a field experiment consisting of eight experimental sites spread throughout the reserve (Figure 1).

**Figure 1.**
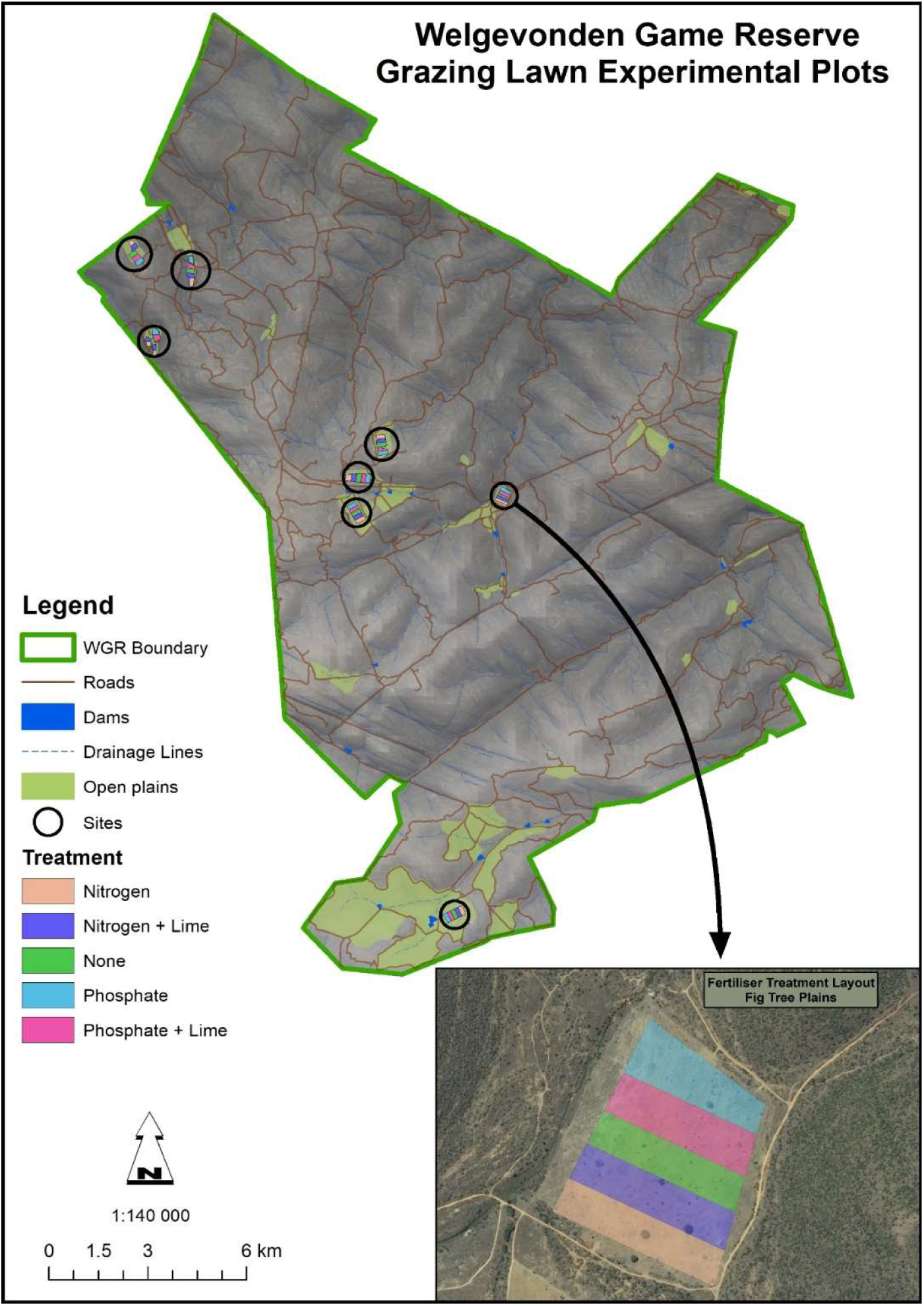
Map of the Welgevonden Game Reserve showing the grazing lawn experimental sites and layout of treatment plots. WGR = Welgevonden Game Reserve

Each site had to be minimally 22.5 hectares to enable testing at a meaningful scale for large herbivores. Due to the mountainous, dissected landscape, the reserve size and form did not allow for more sites. We divided each of the sites into five treatment plots, each minimally 300 m x 150 m (approximately 45,000 m² or 4.5 hectares). The treatment plots had to be large as we wanted to test whether we could create grazing lawns at a landscape scale, and due to the high lion (*Panthera leo*) and leopard (*Panthera pardus*) density, the plots had to be large enough to create relative safety for the grazing herbivores. Exclusion cages were placed in the centre of each treatment plot to allow for an area of comparison in which grazing could not occur (explained in the section on Grass samples). The plots were treated annually between the months of January and February, for three consecutive years, with either the addition of nitrogen (N at 0.25 ton/ha = 25 g/m^2^) or phosphorus (P at 0.50 ton/ha = 50 g/m^2^). The quantities of N and P were chosen based on acceptable levels of these nutrients required for optimum agricultural growth rates. It is difficult for plants to absorb nutrients in soil with a low pH (Chimdi et al. 2012; Higgins et al. 2012); we therefore further investigated the supplementation of lime (Ca) at a concentration of 3 tons/ha (in the form of calcitic lime at 1.5 tons/ha and dolomitic lime at 1.5 tons/ha), to increase the pH of the soil. For that reason, we fertilised plots with nitrogen and lime (N at 0.25 tons/ha and Ca at 3 tons/ha) (calcitic and dolomitic lime) and phosphorus and lime (P at 0.50 tons/ha and Ca at 3 tons/ha) (calcitic and dolomitic lime). We fertilised with N and P separately because we did not intend to increase plant biomass production, but we wanted to increase food quality as to stimulate grazing for sward induction. We included plots with no treatment (controls) to establish if there would be significant differences if there was zero fertilisation (Figure 1).

For the fertilisation of N, we used an ammoniacal nitrate-based fertiliser, SASOL with Limestone Ammonium Nitrate (LAN) and Calcium Ammonium Nitrate (CAN). The fertiliser contains 28% nitrogen of which 50% as ammonium-N and 50% as nitrate-N. In addition, it contains 20% Dolomitic lime resulting in about 4.1% Ca and 2.1% Mg in the product. The P fertiliser was a Yara super phosphate 10.5% P fertiliser (containing P at 105 g per kilogram, S at 106 g per kilogram and Ca at 202 g per kilogram) and both were applied in the form of small granular pellets via a tractor and spreader. Annually, before the plots were treated the encroachment of trees and shrubs was controlled (if required) on all experimental sites (with the exception of in the exclusion cages) by mowing with a tractor and slasher. This reduced the tall forbs and woody species component such as *Sida cordifolia*, *Dichrostachys cinerea* and other *Vachellia spp*. to a height of 10cm above the ground (Van der Waal 2010). Woody species were not desired in the experimental plots because they impede food uptake by herbivores, can conceal large predators such as lion, have different uptake characteristics of nutrients (cf. Van der Waal 2010) and can shade out lawn grass species affecting growth.

### Soil samples

Over the three-year period 2016-2018, we collected a total of 1290 soil samples during the first (January to April), second (May to August) and third periods (September to December) of each of the three years. We sampled these three periods to account for the possible variation in soil mineral status over the year, for instance due to fluctuations in mineralization rates after droughts or rain. These samples were collected from the centre of each of the 40 treatment plots, at three depths, 0-5, 10-15 and 25-30cm, including the control plots. We collected the soil using a narrow soil auger. The reason for collecting the three different depths was to establish leaching levels of the various minerals after rain.

Prior to the analyses, we dried the samples in a forced-air oven at a temperature of 40°C (low temperature to prevent N-losses) till dry and sieved them through a 0.5-1 mm mesh sieve. Soil sample minerals were analysed either as a percentage in the case of acid saturation, milligrams per kilogram (mg/kg) for P, K, Na, Ca, Mg and N, grams per centimetre cubed (g/cm³) for density and for pH the KCl extractable was used to remove the aluminium. pH samples were mixed with 1.0 molar potassium chloride at a ratio of 1: 2.5 and read with a pH meter and a glass electrode. P was extracted using BRAY I extractant and determined colorimetric. K, Na and Ca samples were extracted by 1.0 molar ammonium acetate and determined by inductively coupled plasma spectroscopy (Barnard et al. 1990; Rayment and Lyons 2011; NviroTek Laboratories 2018).

### Grass samples

To test the effect of moderate to heavy grazing by herbivores, we set up grazing exclusion cages from January 2017 (1 m x 1 m; 1.2 m high) in the centre of each treatment plot where *Cynodon dactylon,* the most prevalent lawn grass species, was in abundance. Grazing exclusion cages were set up to establish the original and change in grass species composition and biomass in the presence and absence of herbivores over time. The cages remained in the same locations throughout the two-year period to ensure that grazing never became a factor in the change of grass composition in the cages overtime. Over the two-year period 2017-2018, we collected a total of 1262 grass samples from both inside (636) and outside (626) the exclusion cages during the first (January to April), second (May to August) and third periods (September to December) of the year (if grass was present). The grass leaves were cut in the 1 m x 1 m exclusion cages and a 1 m x 1 m patch adjacent to the exclusion cages, within a radius of 10 m of the exclusion cage. The samples were cut from a height of 10cm above ground level. We included all the fresh leaf material and dry leaf material (which was available to be grazed), which varied depending on the time of year the grass samples were collected. The grass samples from each individual treatment plot were sorted and identified at species level using the Welgevonden grass species reference list (Appendix A). The mass of the different grass species was measured and recorded to assess the individual grass species abundance. In the field, before each grass sample was collected, the biomass of the grass leaf was measured both in the exclusion cages and adjacent to the exclusion cages within the 1m x 1m patch. The comparable measurements were taken using a calibrated disc pasture meter (DPM) to compare the biomass of the grass leaf outside and inside the cage within each treatment plot. The DPM used has a diameter of 46cm and weighs 1.34kg, with the disk being dropped from a height of 1.7m above ground level. The calibration equation used was the mean of five previously used formulae and was Y = biomass in kg/ha and X = grass height in cm.

To analyse the grass samples mineral content, we air-dried the combined grass leaf samples from each individual treatment plot, followed by drying in an oven at 70°C. We then analysed the grass sample minerals, either as a percentage in the case of N, Ca, Mg, K, S and P or mg/kg as with Na, Fe, Mn, Cu, Zn and B. N was determined by a Dumas (total combustion) method and measured against organic material with a known N content. All the other compounds were digested with a Nitric and Perchloric acid mixture at 230°C, allowing for the elements to be determined by Inductive Coupled plasma (NviroTek Laboratories 2018; AGRILASA 2019). Drying and chemical analyses were carried out at NviroTek laboratories, which is an independent certified analytical testing facility. Nvirotek uses the international accredited determination of total nitrogen by the DUMAS combustion method as approved by the Agricultural laboratory Association of Southern Africa.

### Environmental impact of fertilisation

To ascertain that our large-scale fertilisation had no negative impact on the water system or water quality close to the grazing lawn experimental sites, Welgevonden Game Reserve contracted three independent laboratories (NviroTek Laboratory, Hartbeespoort, South Africa; UIS Analytical services, Centurion, South Africa and the Witwatersrand University, Johannesburg, South Africa) to take and analyse separate samples from surface water close to the experimentally fertilised plots. The samples were collected from one site, *viz*. ‘Figtree Plains’, as this is the only site which has a river system in its immediate vicinity. Samples were taken in October 2017, April 2018 and August 2018, with each sample being collected at three different locations within the river. Sample 1 taken upstream from the site (24°17’ 24.21” S 27° 49’ 42.73” E), sample 2 taken just below the site (24°17’ 07.71” S 27° 49’ 53.57” E) and sample 3 taken a few kilometres downstream from the site (24°14’ 47.53” S 27° 50’ 43.13” E). The analyses showed that there was no measurable change in the nitrate levels at any of the test locations. Concentrations of N (<0.5 mg/l) and P (<5 mg/l) showed that the system is extremely oligotrophic and in an unspoiled condition. There was no discernible effect of the fertilisation on the river system or water quality.

### Statistical analysis

We tested our hypothesis using linear mixed models (LMMs) for each of the minerals in the soil and in the grass leaf, and the fraction of lawn grass in the total grass biomass as response variables, the fertilisation treatments as a fixed factor, and year, sampling moment and site as random factors. The differences between the fertilisation treatments were compared using Šidák multiple comparisons tests. We also ran LMMs to test the correlation between N, P and Ca concentrations in the soil and in the grass leaf. Again, site and year were added as random factors. We used year as a random factor as we were not interested in differences between the years assuming that the response of the grasses to the mineral addition was within a few weeks after the application (Van der Waal et al. 2011a). We checked whether the residuals of the LMMs were normally distributed and, if not, log_e_ transformation of the response variable was used. For the fraction of lawn grass in the total grass biomass, we used the arcsine square root transformation. Analysis were performed in SPSS v. 23 (SPSS Inc., Chicago, USA) and Canoco 5.12 (Ter Braak and Smilauer 1998).

## Results

### Soil minerals

In the soil samples that we collected at different depths (0-5, 10-15 and 25-30cm), there were no major differences between the treatment plots in the sample levels from 10-15 and 25-30cm (Appendix B). The three random factors, year, sampling moment and site were not significant. This shows that the increased mineral availability due to fertilisation was mainly found in the upper soil layer from 0-5cm, suggesting that there was no leaching during the period of research, and that minerals that had been supplied were available for the vegetation to utilise during our measurement period. Hence, we only considered the mineral quantities in the upper soil layer in further analyses. We found differences in levels of P, N, Ca and Mg in the 0-5cm soil layer between the treatments (Figure 2, Table 1). The N and P values in the soil were 2 to 3 times higher in the plots fertilised with N, P and the combination of Ca, than in the control plots. As expected, there were higher concentrations of N in the soil of plots which had been treated with N, and higher P concentrations in those that were fertilised with P, which concurs with the findings of Higgins et al. (2012). Ca and Mg concentrations were higher in the soils of plots treated with lime.

**Figure 2.**
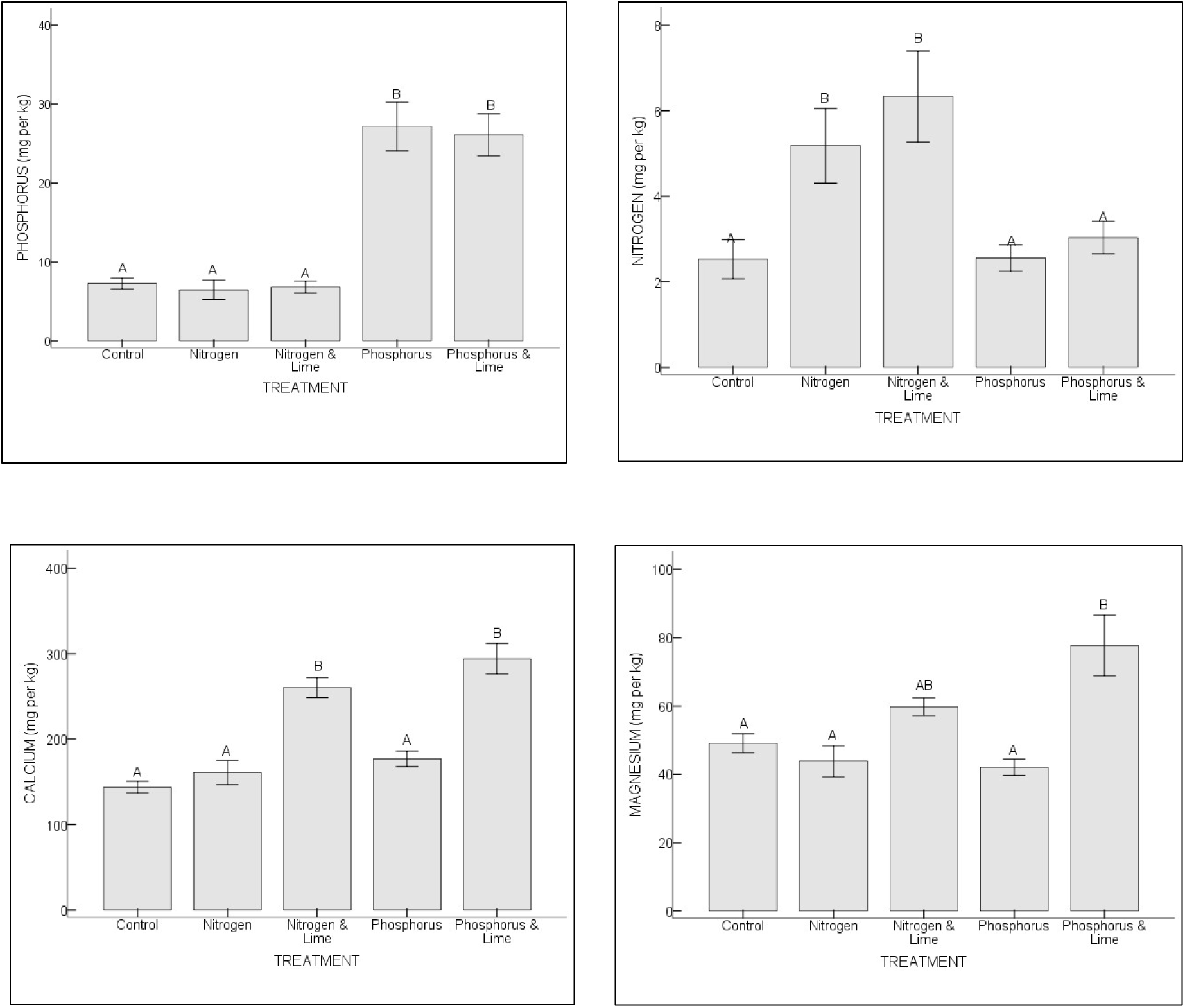
Soil minerals (phosphorus, nitrogen, calcium, magnesium – all expressed in mg/kg) in the 0-5cm soil layer per treatment (control, nitrogen, nitrogen and calcitic and dolomitic lime, phosphorus, phosphorus and calcitic and dolomitic lime). Error bars represent the standard error of the mean. Letters indicate significant differences between the treatments based on Šidák multiple comparisons tests using a linear mixed model (see Table 1 for the statistics)

**Table 1.** Results of the linear mixed model for differences in soil nutrients in the 0-5cm soil layer between the treatments. Separate mixed effects models were run for each mineral (see Figure 2 for the graphs; the differences between the treatments are indicated in the figure using letters). Year and Site were the random factors, estimation method was REML and the sample size n = 361

|  | Statistic | P | N | Ca | Mg |
| --- | --- | --- | --- | --- | --- |
| Treatment | F | 12.4 | 3.4 | 8.2 | 3.5 |
|  | df | 34.7 | 34.9 | 35.0 | 35.0 |
|  | P | <0.001 | <0.05 | <0.001 | <0.05 |
| Year | Wald Z | 1.0 | 0.9 | 0.9 | 0.5 |
|  | P | 0.338 | 0.358 | 0.362 | 0.617 |
| Site | Wald Z | 2.9 | 2.3 | 3.3 | 2.8 |
|  | P | <0.01 | <0.05 | <0.001 | <0.01 |

### Grass minerals

The treatments resulted in differences in concentrations of N, P, Ca, Mg, S and Mn (Figure 3, Table 2) in the plants. As per our first hypothesis N, P and Ca in the grass leaf were at higher values in the plots treated with these minerals. The three random factors, year, sampling moment and site were not significant.

**Figure 3.**
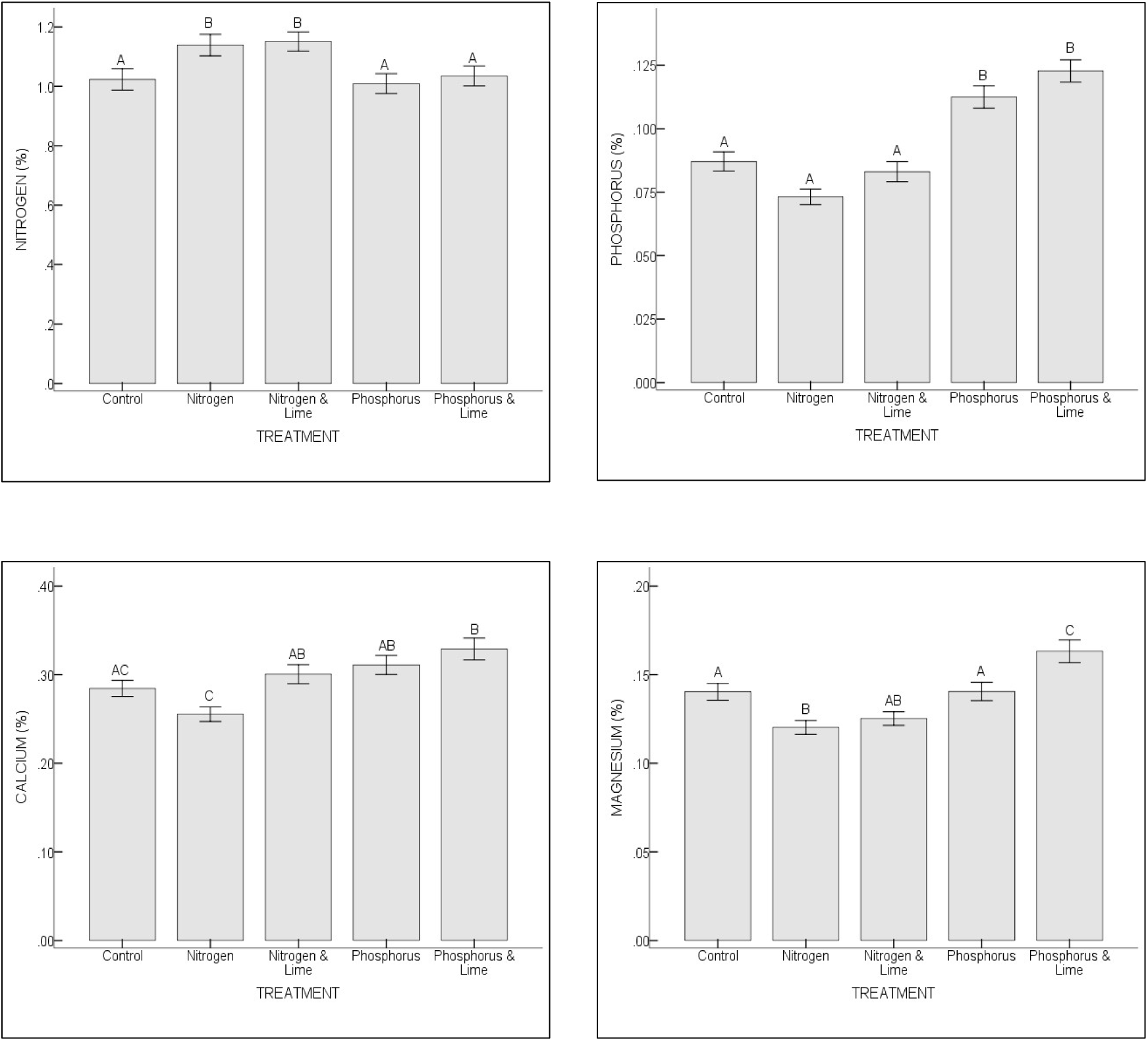
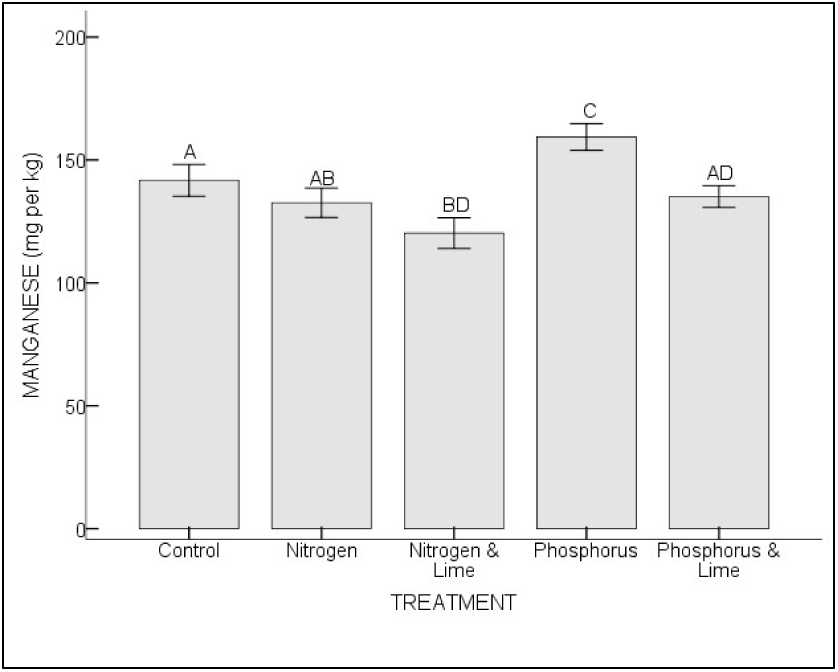
Grass minerals (nitrogen, phosphorus, calcium, magnesium, manganese (Mn) – all expressed as a percentage except for Mn which is expressed in mg/kg) with significant values per treatment (control, nitrogen, nitrogen and calcitic and dolomitic lime, phosphorus, phosphorus and calcitic and dolomitic lime). Error bars represent the standard error of the mean. Letters indicate significant differences between the treatments based on Šidák multiple comparisons tests using a linear mixed model (see Table 2 for the statistics)

**Table 2.** Results of the linear mixed model for differences in nutrients in the grass between the treatments (see Figure 3 for the graphs; the differences between the treatments are indicated in the figure using letters). Year and Site were the random factors, estimation method was REML and the sample size n = 569. Separate mixed effects models were run for each mineral

|  | Statistic | N | P | Ca | Mg | S | Mn |
| --- | --- | --- | --- | --- | --- | --- | --- |
| Treatment | F | 4.1 | 36.5 | 7.8 | 12.9 | 7.1 | 8.3 |
|  | df | 557.0 | 556.1 | 556.1 | 556.1 | 556.1 | 557.0 |
|  | P | <0.01 | <0.001 | <0.001 | <0.001 | <0.001 | <0.001 |
| Year | Wald Z | 0.3 | 0.7 | 0.7 | 0.6 | 0.70 | - |
|  | P | 0.760 | 0.486 | 0.507 | 0.545 | 0.482 | - |
| Site | Wald Z | 1.2 | 1.8 | 1.7 | 1.7 | 1.8 | 1.8 |
|  | P | 0.257 | <0.1 | <0.1 | <0.1 | <0.1 | <0.1 |

Unlike what we had hypothesized we found that the correlation between N in the soil and N in the grass leaf was negligible and not significantly higher than the control plots. However, there was a positive correlation between P in the soil and the P in the grass leaf, as well as for Ca in the soil and Ca in the grass leaf (Figure 4 and Table 3). The P value in the grass leaf of the plots fertilised with P and/or Ca was 1.5 times higher than in the grass leaf in the control plots.

**Figure 4.**
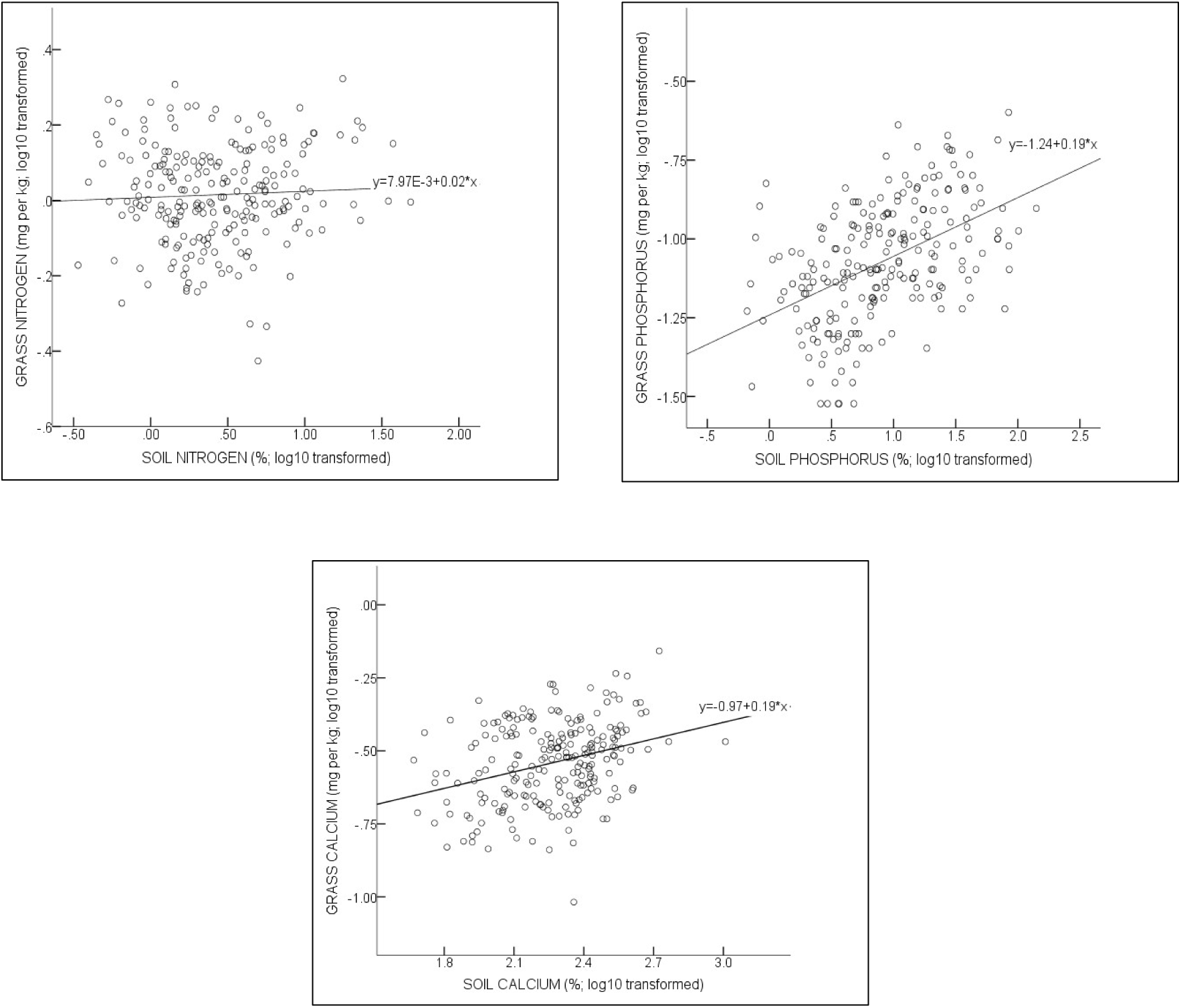
Correlation between nitrogen, phosphorus and calcium in the soil (all expressed in mg/kg; log_10_ transformed) and nitrogen, phosphorus and calcium in the grass leaf (all expressed as a percentage; log_10_ transformed) (see Table 3 for the statistics)

**Table 3.**
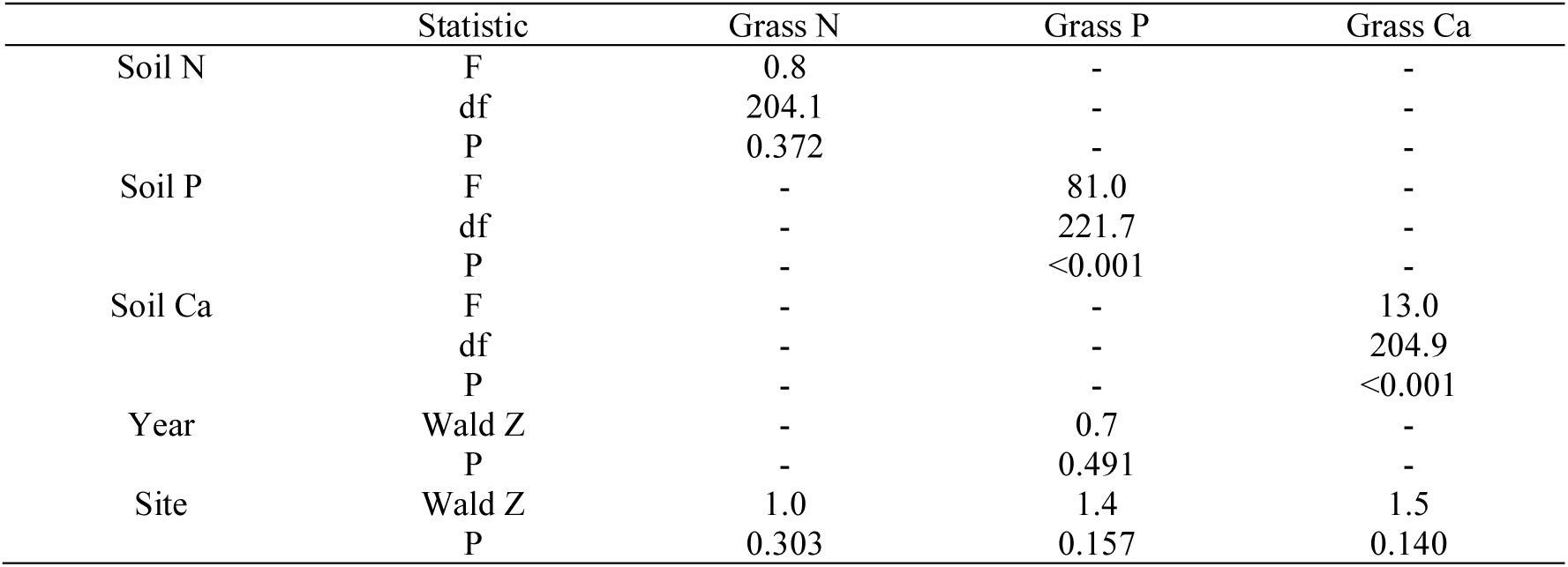
Results of the linear mixed model for the correlation between (i) N, (ii) P and (iii) Ca in the soil and the grass (see Figure 4 for the graphs). Year and Site were the random factors, estimation method was REML, and the sample size n = 224. All variables were log10 transformed

|  | Statistic | Grass N | Grass P | Grass Ca |
| --- | --- | --- | --- | --- |
| Soil N | F | 0.8 | - | - |
|  | df | 204.1 | - | - |
|  | P | 0.372 | - | - |
| Soil P | F | - | 81.0 | - |
|  | df | - | 221.7 | - |
|  | P | - | <0.001 | - |
| Soil Ca | F | - | - | 13.0 |
|  | df | - | - | 204.9 |
|  | P | - | - | <0.001 |
| Year | Wald Z | - | 0.7 | - |
|  | P | - | 0.491 | - |
| Site | Wald Z | 1.0 | 1.4 | 1.5 |
|  | P | 0.303 | 0.157 | 0.140 |

### Change in composition of grasses

In line with our second hypothesis, the fraction of *Cynodon dactylon* outside the cages was higher (with an increased biomass) than inside for all the plots except for the treatment plots of P with lime, where the difference was not statistically significant (Figure 5, Table 4). Overall, we found a larger increase in biomass of lawn grasses (*Cynodon spp*.) in the 1m x 1m patches outside the exclusion cages than inside. The three random factors, year, sampling moment and site were not significant. We found a higher proportion of *Cynodon spp*. within the treatment plots than within the control plots with no nutrients added, with the exception of P treatment plots, which did not have significantly different results to the control plot.

**Figure 5.**
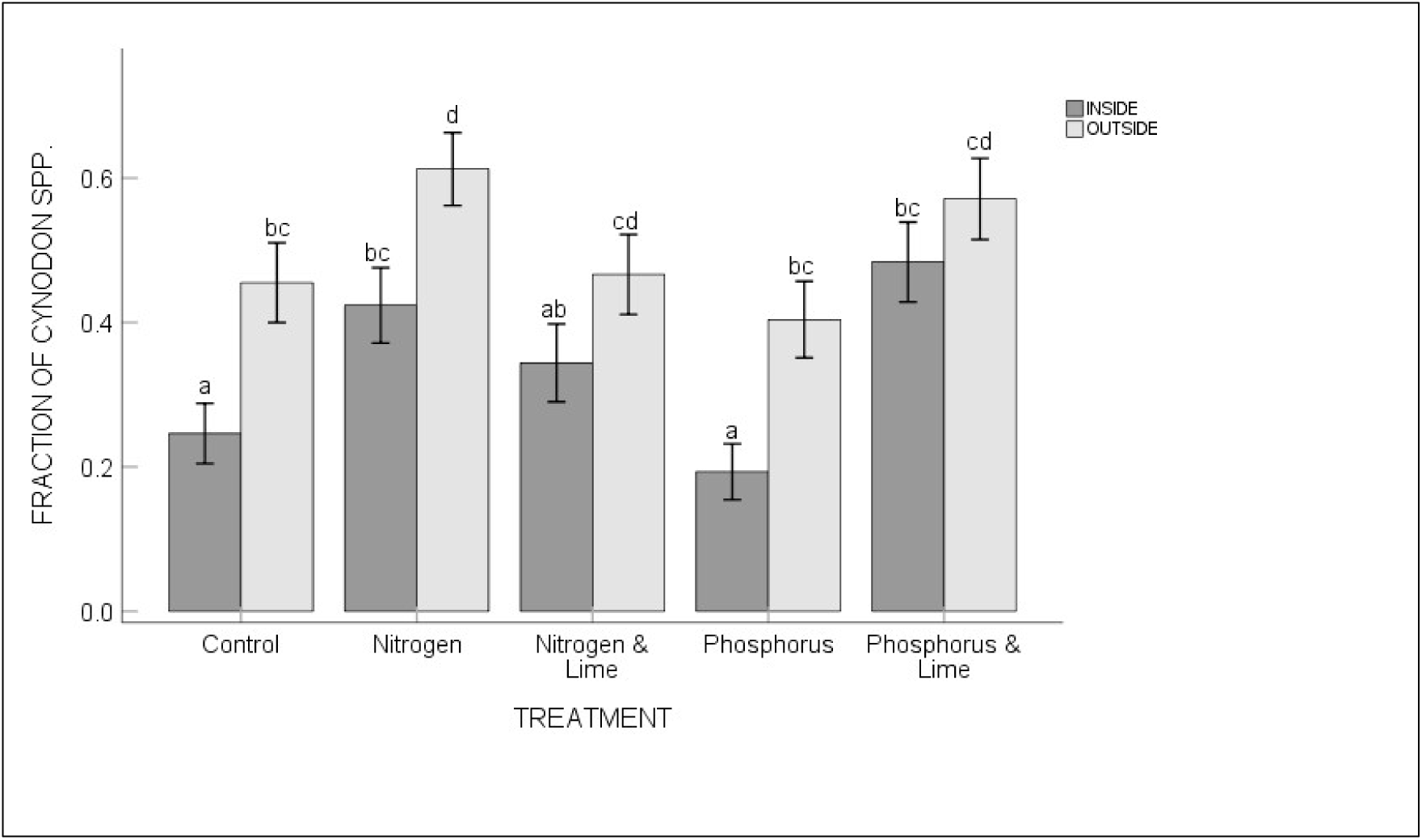
The proportion of *Cynodon dactylon* (biomass Cynodon / total biomass) inside and outside the cages per treatment (control, nitrogen, nitrogen and calcitic and dolomitic lime, phosphorus, phosphorus and calcitic and dolomitic lime). Error bars represent the standard error of the mean. Letters indicate significant differences between the treatments based on Šidák multiple comparisons tests using a linear mixed model (see Table 4 for the statistics)

**Table 4.** Results of the linear mixed model for differences in the fraction of *Cynodon dactylon* (biomass *Cynodon* / total biomass) between the treatments and inside or outside the cages (see Figure 5 for the graphs). Year, Period and Site were the random factors, estimation method was REML

|  |  |  |
| --- | --- | --- |
| Treatment | F | 11.7 |
|  | df | 4 |
|  | P | < 0.001 |
| Inside/Outside | F | 45.9 |
|  | df | 1 |
|  | P | < 0.001 |
| Treatment x Inside/Outside | F | 0.8 |
|  | df | 4 |
|  | P | 0.527 |
| Year | Wald Z | - |
|  | P | - |
| Period | Wald Z | 0.9 |
|  | P | 0.342 |
| Site | Wald Z | 1.9 |
|  | P | <0.1 |
| n |  | 628 |

## Discussion

We tested the hypothesis that the addition of minerals to a nutrient poor savanna increases mineral levels in the soil and in the grass leaf, resulting in an increase in lawn grass proportions. There are about one hundred lawn grass species in South Africa (native and introduced) that are stoloniferous (Fish et al. 2015). Most of these stoloniferous species can be induced through grazing or mowing to become lawn-forming. Our results support this hypothesis as there was a significant difference between the proportions of lawn grasses inside the cages, which could not be grazed versus outside the cages, which could be grazed freely by herbivores. Seymour et al. (2013) and Veldhuis et al. (2014) state that grazing lawns are generally found in areas with relatively high soil nutrient status. However, Cromsigt and Ollf 2008, who experimented on plots 8m x 8m in size, showed that plots which were unfertilised, did not develop grazing lawns. Our experiments were undertaken on a far larger spacial scale with rigorous levels of repetition and showed that increased grazing pressure on the fertilised plots stimulated the establishment of grazing lawns on the unfertilised plots situated between them. Our experiments demonstrate that fertilisation in nutrient poor savannas can trigger the establishment of grazing lawns, however grazing significantly increased the capacity of establishing a grazing lawn without regular mowing of the grass. The results of our study were obtained under the condition that initial mowing is required to enable the grazers to graze by removing stalks and woody species that hinder their foraging (Drescher et al. 2006). The majority of research done on the addition of nutrients were undertaken with combinations of N and P, where our experiments were novel in the fact that we wanted to establish outcomes of the ability of grazing lawn formation on separate plots emphasising the different outcomes after the fertilisation with N and P separately. We found that the addition of P and lime (calcitic and dolomitic) significantly increased the amounts of available P and lime (Ca) in the grass leaf, which was also found in a study by Higgins et al. (2012). Because our study area is dominated by soils with a low pH, adding calcitic and dolomitic lime increases the pH thus promoting P-availability in the soil to the plants (Chimdi et al. 2012; Higgins et al. 2012). N did not show significant increased mineral levels in the grass leaf , this is likely due to induced rapid growth, with the consequent dilution of minerals (Schnyder et al. 2000; Gastal and Lemaire 2015). Grass species were not separated prior to the analysis of the nutrients, thus we cannot ascertain if the response is due to increased nutrient uptake by the grasses, or a switch in species composition to grasses with higher intrinsic nutrient concentrations. Hempson et al. 2014 show that the impact of repeated grazing, results in a switch from tall to short grass, with the short grass actively growing and having a higher nutrient content than the tall grass. However, we show that there is an improvement of the forage quality, whether enhanced through nutrient uptake, a change towards a more nutrient rich species composition or a grazing induced change in structure of the grass. The differences in the effect of N and P fertilisation on the minerals in the grass was also found by Van der Waal et al. (2011a), who used similar levels of fertilisation. They showed that P fertilisation increased leaf P concentrations more in grasses than trees, whereas N fertilisation increased leaf N concentration moderately in both trees and grasses. We conclude from this study that N and more especially P increased in the soil when N or P were applied, but that P was the mineral which was made available to grazers in the grass leaf. The data suggests that P is limiting in this dystrophic savanna type (Figure 4). This is supported by the differences in grass biomass in the absence of grazing (inside the cages) between the N and P plots. We found that the total grass biomass in the exclusion cages on treatment plots where P was added was higher than in the treatment plots where N was added (LMM for 2017: P = 0.012, F_1,59.0_ = 6.7, but not for 2018: P = 0.823, F_1,43.7_ = 0.05). The growth of lawn grasses (especially *Cynodon dactylon*) is likely due to the increased availability of minerals in the soil for growth and stolon formation, which increased the proportions of lawn grass cover in and outside of the cages. Furthermore, the increase in mineral availability in the soil and grass leaf attracts more grazers to the fertilised areas, which ensures that the grazing lawns remain well-grazed (Schroder 2021), resulting in the long-term maintenance of the lawn grass patches. Our results also show that when the grass is not grazed (inside the cages), lawn grasses (especially couch grass) are more readily outcompeted by other grass species (Figure 5). The control plots, which did not have any nutrients added to them, showed a significant difference in the proportion of lawn grasses between the grazed (outside the cage) and not grazed (inside the cage). We posit that this is due to the fact that at each site, the control plot was situated between the fertilised plots (Figure 1), suggesting that the herbivores grazed the plots ‘by default’ as they moved between the various treatment plots, as shown in Schroder (2021).

Fire reduces grazing lawn success indirectly by creating green flushes in other areas which draw the grazers away from the lawns (Bond and Archibald 2003; Van Langevelde et al. 2003) and thus restricting the development and maintenance of lawns and perpetuating fire-driven systems. Increases in mineral concentration in the grass may result in grazing lawns that are herbivore-driven (Waldram et al. 2007; Archibald 2008; Cromsigt and Olff 2008; Novellie and Gaylard 2013; Hempson et al. 2014; Zwerts et al. 2015). In nutrient poor savannas with grasses of low nutritional value, herbivores may have poor body condition resulting in low levels of nutritional value in their milk and thus a negative effect on fecundity. This has been reflected by low P and N concentrations in faeces in several studies (Dörgeloh et al. 1998; Grant et al. 2000). To ensure high numbers of herbivores in nutrient poor savannas (for instance desired for suppressing dangerous wild fires or good viewing potential for the tourism industry), the forage quality of the grass needs to be improved and maintained.

In mesic savannas (rainfall > 650 mm), such as Kruger National Park, where there is a higher amount of nutrients available, high grazing pressure may lead to the replacement of palatable species by unpalatable species (Grant et al. 2011). Our study area has a historic annual rainfall of > 650 mm but is situated on soils with much poorer nutrient availability than found nearly anywhere in the Kruger National Park, even on the granitic sandy soils, and much poorer than the volcanic soils there (Ferwerda et al. 2006; Skidmore et al. 2009; Ramoelo et al. 2013). Figure 5 shows how grazing pressure and manipulation outside the cages increased the proportions of the lawn grasses over a period of time and changed the grass species from tall nutritionally poor to short nutritionally rich grasses.

## Practical implications

With these findings we contribute to the understanding of under what conditions grazing lawns can establish or be established. The manipulation of grazing lawns through the addition of P and Ca specifically can be attributed to the increased mineral value of grass leaf in nutrient poor savannas. Future studies should test the hypothesis that herbivores will maintain grazing lawns in this state in nutrient poor savannas such as Welgevonden Game Reserve. It is critical to continue monitoring the responses of the treatments in the long-term, especially if the application of fertiliser is stopped. Several studies of N addition in savanna ecosystems have been undertaken, but there are many pathways for N loss and dilution. In contrast, there have been only a few studies pertaining to the role of P in grazing lawn formation, and P is often highly conserved in nutrient-poor ecosystems. It has been suggested that P fertilization treatments will have substantially longer lasting effects on lawn persistence, and could eventually lead to N accumulation via herbivore attraction, which needs monitoring. Grazing lawns can stop or prevent wild fires (which can be devastating to property and upset management schemes), and are key to a successful game viewing industry (eco-tourism). It is difficult to make eco-tourism financially viable in nutrient poor areas, where it was previously shown that agricultural farming was unproductive and financially unsuccessful.

## Supporting information

Supporting Documentation A

Supporting Documentation B

---

**Appendix A.** Welgevonden Game Reserve grass species list (Jonathan Swart – Reserve Ecologist - Personal communication) with the grazing values obtained from the guide to grasses of Southern Africa (Van Oudtshoorn 2002)

| Scientific name | English name | Grazing value |
| --- | --- | --- |
| *Alloteropsis semialata* | Black-seed Grass | Average |
| *Andropogon appendiculatus* | Vlei Bluestem | High |
| *Andropogon chinensis* | Hairy Blue Grass | Average |
| *Andropogon eucomus* | Snowflake Grass | Low |
| *Andropogon huillensis* | Large Silver Andropogon | Average |
| *Andropogon schirensis* | Stab Grass | Average |
| *Aristida adscensionis* | Annual Three-awn | Low |
| *Aristida congesta* ssp. *barbicollis* | Spreading Three-awn | Low |
| *Aristida congesta* subsp. *congesta* | Tassel Three-awn | Low |
| *Aristida diffusa burkei* | Iron Grass | Low |
| *Aristida junciformis* ssp. *junciformis* | Ngongoni Three-awn | Low |
| *Aristida meridionalis* | Giant Three-awn | Low |
| *Aristida scabrivalvis* ssp. *scabrivalvis* | Purple Three-awn | Low |
| *Aristida stipitata* ssp. *graciliflora* | Long-awned-three-awn | Low |
| *Aristida transvaalensis* | Rock Three-awn | Low |
| *Bewsia biflora* | False Love Grass | Low |
| *Brachiaria brizantha* | Common Signal Grass | Average |
| *Brachiaria deflexa* | False Panicum | Average |
| *Brachiaria eruciformis* | Sweet Signal Grass | Average |
| *Brachiaria nigropedata* | Black-footed Signal Grass | High |
| *Brachiaria serrata* | Velvet Signal Grass | Average |
| *Cenchrus ciliaris* | Blue Buffalo Grass | High |
| *Chloris gayana* | Rhodes Grass | Low |
| *Chloris pycnothrix* | Spiderweb Grass | Low |
| *Chloris virgata* | Feathered Chloris | Average |
| *Chrysopogon serrulatus* | Golden Beard Grass | High |
| *Cymbopogon caesius* | Broad-leaved Turpentine Grass | Low |
| *Cymbopogon pospischilii* | Narrow-leaved Turpentine Grass | Low |
| *Cynodon dactylon* | Couch Grass | High |
| *Cynodon nlemfuensis* | Star Grass | High |
| *Dactyloctenium aegyptium* | Common Crowfoot | Average |
| *Dactyloctenium giganteum* | Giant Crowfoot | High |
| *Digitaria brazzae* | Brown Finger Grass | High |
| *Digitaria eriantha* | Finger Grass | High |
| *Digitaria longiflora* | False Couch Grass | Average |
| *Digitaria monodactyla* | One-finger Grass | Average |
| *Digitaria ternata* | Black-seed Finger Grass | Low |
| *Digitaria velutina* | Flaccid Finger Grass | Low |
| *Diheteropogon amplexans* | Broad-leaved Bluestem | Average |
| *Diplachne fusca* | Swamp Grass | High |
| *Echinochloa colona* | Jungle Rice | High |
| *Eleusine coracana* ssp. *africana* | Goose Grass | Low |
| *Elionurus muticus* | Wire Grass | Low |
| *Enneapogon pretoriensis* | Nineawn Grass | Low |
| *Eragrostis aspera* | Rough Love Grass | Low |
| *Eragrostis biflora* | Shade Eragrostis | Low |
| *Eragrostis caesia* | Weeping Lovegrass | Low |
| *Eragrostis capensis* | Heart-seed Love Grass | Average |
| *Eragrostis chloromelas* | Curly Leaf | Average |
| *Eragrostis curvula* | Weeping Love Grass | High |
| *Eragrostis gummiflua* | Gum Grass | Low |
| *Eragrostis heteromera* | Bronze Love Grass | Average |
| *Eragrostis lehmanniana* | Lehmann's Love Grass | Average |
| *Eragrostis micrantha* | Finesse Grass | Low |
| *Eragrostis nindensis* | Wether Love Grass | Average |
| *Eragrostis patens* | Japanese Lovegrass | Low |
| *Eragrostis pallens* | Broom Love Grass | Low |
| *Eragrostis plana* | Tough Love Grass | Low |
| *Eragrostis pseudosclerantha* | Footpath Love Grass | Low |
| *Eragrostis racemosa* | Narrow Heard Love Grass | Average |
| *Eragrostis rotifer* | Pearly Love Grass | Average |
| *Eragrostis superba* | Sawtooth Love Grass | Average |
| *Eragrostis viscosa* | Sticky Love Grass | Low |
| *Eustachys paspaloides* | Fan Grass | High |
| *Fingerhutia africana* | Thimble Grass | Average |
| *Heteropogon contortus* | Spear Grass | Average |
| *Hyperthelia dissoluta* | Yellow Thatching Grass | Average |
| *Hyparrhenia filipendula* | Fine Thatching Grass | Average |
| *Hyparrhenia hirta* | Common Thatching Grass | Average |
| *Imperata cylindrica* | Cottonwool Grass | Low |
| *Ischaemum fasciculatum* | Hippo Grass | Average |
| *Leersia hexandra* | Southern Cutgrass | High |
| *Leptocarydion vulpiastrum* | Spade Grass | Low |
| *Loudetia simplex* | Common Russet Grass | Average |
| *Melinis nerviglumis* | Bristle-leaved Red Top | Average |
| *Melinis repens* ssp. *repens* | Natal Red Top | Low |
| *Miscanthus junceus* | Wireleaf Daba Grass | Average |
| *Monocymbium ceresiiforme* | Boat Grass | Average |
| *Panicum coloratum* var. *coloratum* | White Buffalo Grass | High |
| *Panicum deustum* | Broad-leaved Panicum | High |
| *Panicum dregeanum* | Plum Panicum | Average |
| *Panicum maximum* | Guinea Grass | High |
| *Panicum natalense* | Natal Panicum | Low |
| *Paspalum dilatatum* | Common Paspalum | High |
| *Paspalum scrobiculatum* | Ricegrass Paspalum | Average |
| *Paspalum urvillei* | Giant Paspalum | Average |
| *Pennisetum macrourum* | Riverbed Grass | Average |
| *Perotis patens* | Cat's Tail | Low |
| *Pogonarthria squarrosa* | Herringbone Grass | Low |
| *Sacciolepis typhura* | Purple-hood Grass | Low |
| *Schizachyrium jeffreysii* | Silky Autumn Grass | Low |
| *Schizachyrium sanguineum* | Red Autumn Grass | Low |
| *Schmidtia pappophoroides* | Sand Quick | High |
| *Setaria lindenbergiana* | Mountain Bristle Grass | High |
| *Setaria pumila* | Shade Bristle Grass | Low |
| *Setaria sphacelata* var. *sphacelata* | Common Bristle Grass | High |
| *Setaria verticillata* | Bur Bristle Grass | High |
| *Sporobolus africanus* | Ratstail Dropseed | Low |
| *Sporobolus festivus* | Red Dropseed | Low |
| *Sporobolus fimbriatus* | Bushveld Dropseed | High |
| *Sporobolus pyramidalis* | Catstail Dropseed | Low |
| *Themeda triandra* | Redgrass | Average |
| *Trachypogon spicatus* | Giant Spear Grass | Low |
| *Tragus berteronianus* | Common Carrot Seed Grass | Low |
| *Tragus racemosus* | Large Carrot Seed Grass | Low |
| *Trichoneura grandiglumis* | Small Rolling Grass | low |
| *Triraphis schinzii* | Sandveld Broom Grass | Low |
| *Tristachya biseriata* | Trident Grass | Average |
| *Tristachya leucothrix* | Hairy Trident Grass | Average |
| *Tristachya rehmannii* | Trident Grass | Low |
| *Urelytrum agropyroides* | Quinine Grass | Low |
| *Urochloa mosambicensis* | Bushveld Signal Grass | High |
| *Urochloa panicoides* | Garden Signal Grass | Low |

**Appendix B.**
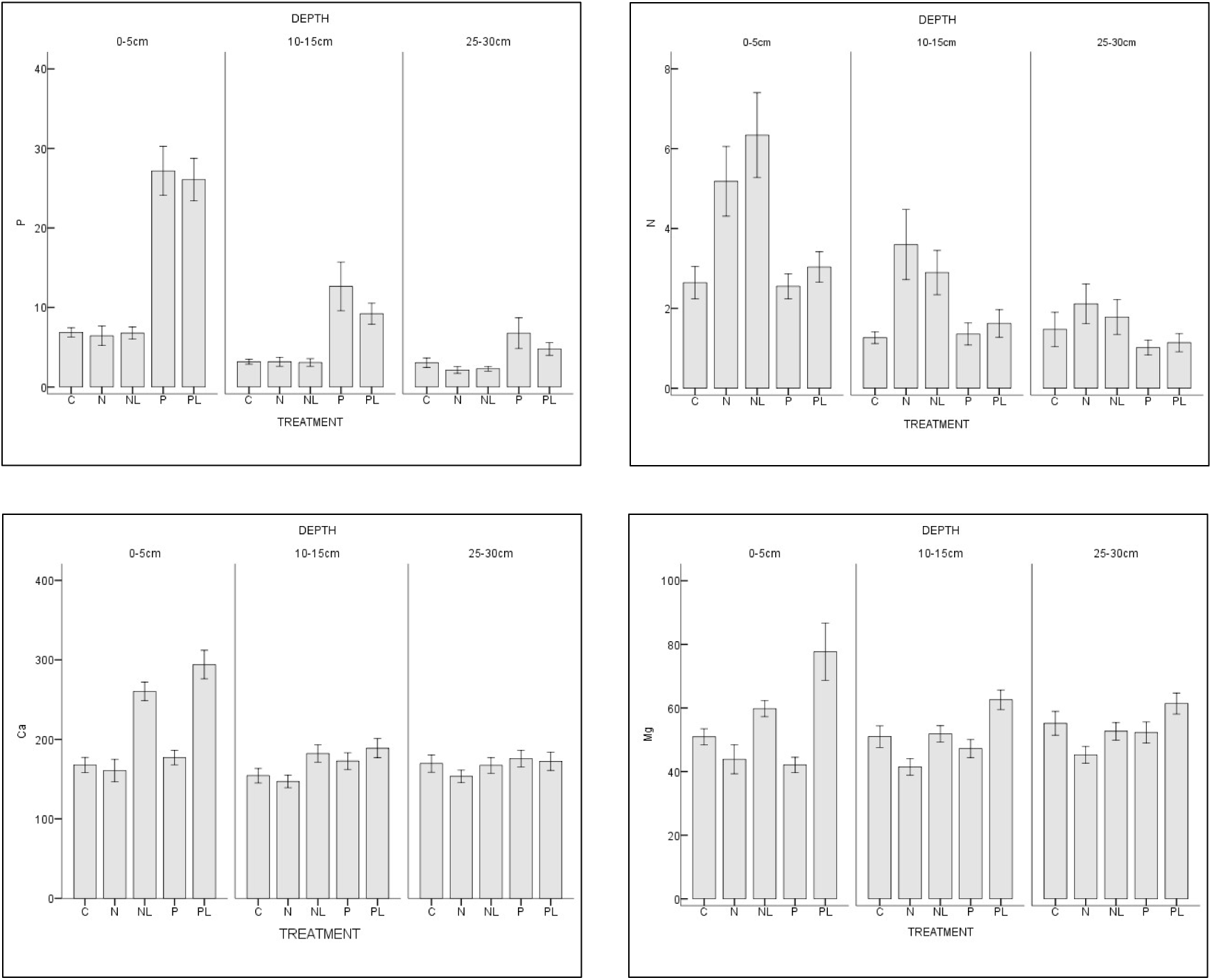
Soil nutrients (P – phosphorus, N – nitrogen, Ca – calcium, Mg – magnesium – all expressed as mg/kg) in the soil layer at 0-5cm, 10-15cm and 25-30cm per treatment (C = control, N = nitrogen, NL = nitrogen and calcitic and dolomitic lime, P = phosphorus, PL = phosphorus and calcitic and dolomitic lime). Error bars represent the standard error of the mean

